# Rapid and efficient C-terminal labeling of nanobodies for DNA-PAINT

**DOI:** 10.1101/389445

**Authors:** Valentin Fabricius, Jonathan Lefèbre, Hylkje Geertsema, Stephen F. Marino, Helge Ewers

## Abstract

Single molecule localization-based approaches to superresolution microscopy (SMLM) create images that resolve features smaller than the diffraction limit of light by rendering them from the sequentially measured positions of thousands of individual molecules. New SMLM approaches based on the transient binding of very bright dyes via DNA-DNA interaction (DNA-PAINT) allow the resolution of dyes only a few nanometers apart *in vitro*. This imaging of cellular structures requires the specific association of dyes to their targets, which results in an additional “linkage error”. This error can be minimized by using extremely small, single-domain antibody-based binders such as nanobodies, but the DNA-oligomers used in DNA-PAINT are of significant size in comparison to nanobodies and may interfere with binding. We have here developed an optimized procedure based on enzymatic labeling and click-chemistry for the coupling of DNA oligomers to the nanobody C-terminus, which is located on the opposite side of the epitope-binding domain. Our approach allows for straightforward labeling, purification and DNA-PAINT imaging. We performed high efficiency labeling of two different nanobodies and show dual color multiplexed SMLM to demonstrate the general applicability of our labeling scheme.

## Introduction

Single molecule localization-based superresolution (SMLM) techniques such as PALM[1], STORM [2] and (*d*)STORM [3] are developing at a rapid pace [4–6] with significant improvements in optical strategies, image processing and analysis techniques. The precision with which a single fluorescent molecule can be localized depends largely on the number of photons detected and, with novel approaches involving bright dyes, molecules that are but 5 nm apart can be resolved [7]. While many aspects of SMLM have now reached a high level of maturity, one of the major challenges remains reducing the size of the fluorescent labels themselves to achieve resolutions on the order of the size of biological molecules. In SMLM, the target species itself is not directly imaged – rather, a fluorescent molecule that is either transiently or permanently associated with the target species is detected., with novel approaches, molecules that are spaced about 5 nm apart can be resolved [7]. While many aspects of SMLM have now reached a high level of maturity, one of the major challenges remains reducing the size of the fluorescent labels themselves to achieve resolutions on the order of the size of biological molecules. In SMLM, the target species itself is not directly imaged – rather, a fluorescent molecule that is either transiently or permanently associated with the target species is detected. The upper resolution limit of the experiment – the precision with which the target species can be localized – is therefore determined by the distance of the detected fluorophore from the center of the target species, referred to as ‘linkage error’.

The problem is readily appreciated when one considers an experiment that can theoretically achieve 40 nm resolution in the localization of a target that is detected via a fluorescently labelled IgG molecule – placing the fluorophore up to 10 nm from the target’s center (even further if the label is on a secondary antibody; consider also the error introduced due to the flexibility of such a fluorescent ‘chain’). Clearly, the smaller the detected label, the better. Recently, small binders called nanobodies have been used to deliver dyes to within a few nm of the target structure [8] and indeed this improves resolution significantly in comparison to traditional immunolabeling [8]. Nanobodies are generalized single domain binding modules derived from the heavy-chain only immunoglobulins produced by camelids [9]. Their combination of an enormous potential binding repertoire, comparable and complementary to that of IgGs, in a robust domain obtainable via low cost and straightforward production has made them powerful tools for basic research and biotechnological applications [10–14]. Because these advantages come in a very small package (~15 kDa), nanobodies are quickly becoming indispensable for super resolution microscopic studies. The small size and compact structure of nanobodies (ca. 2-3 nm diameter, compared to ca. 10 nm for an IgG) makes them very attractive as labels in super resolution experiments – particularly if fluorophores can be site-specifically attached via minimal length linkers. Their compact form also has a positive impact on the potential labeling density that can be achieved with them, particularly for continuous filamentous structures (eg. microtubules), minimizing the steric interference inherent to the use of much larger probes [15].

A recent improvement to both the ease and flexibility of labeling and detection techniques is provided by the ‘DNA points accumulation for imaging in nanoscale topography’ method (DNA-PAINT) (Figure 1), in which the target species, or a specific target detection module, is modified with a short DNA oligonucleotide (the “docking” strand) [16,17]. The position of the labeled species is then detected via addition of a complementary oligonucleotide harbouring a fluorophore (the “imager” strand). The low melting temperature (~25°C) of the short, corresponding duplex ensures a high imager strand k_off_ [17,18] and since the fluorophore can only be localized during the brief period the duplex exists, fluorophores are rapidly exchanged, thus constantly allowing new imager strands with fresh fluorophores to illuminate the target, thereby providing the necessary “blinking” for single-molecule localization microscopy (SMLM) (Figure 1). This technique allows the use of very bright, non-blinking dyes and antifade reagents, leading to significantly brighter localizations and thus higher resolution. DNA-PAINT using nanobodies has been reported for the anti-GFP nanobody, but the labeling was based on poorly selective succinimidylester chemistry combined with click-chemistry[19]. As shown in Figure 1f, the DNA-oligomer required for DNA-PAINT is of significant size in comparison with the nanobody and thus coupling directly to the C-terminus as the site furthest away from the epitope-binding domain would be highly desirable. Here we report C-terminal labeling of anti-tubulin and anti-GFP nanobodies for DNA-PAINT. We have developed a streamlined procedure for the site-specific labeling of nanobodies based on the Sortase A (SrtA) reaction [20] and the copper-free strain promoted alkyne-azide cycloaddition (SPAAC) [21] that improves both the ease of handling and the orthogonality of previously reported labeling schemes. The optimized procedure enables a great flexibility of label choice, allowing for multiplexed DNA-PAINT by coupling different oligo binder sequences to different nanobodies. Given their broad applicability we expect these contributions will substantially augment the current super resolution toolbox.

**Figure 1.**
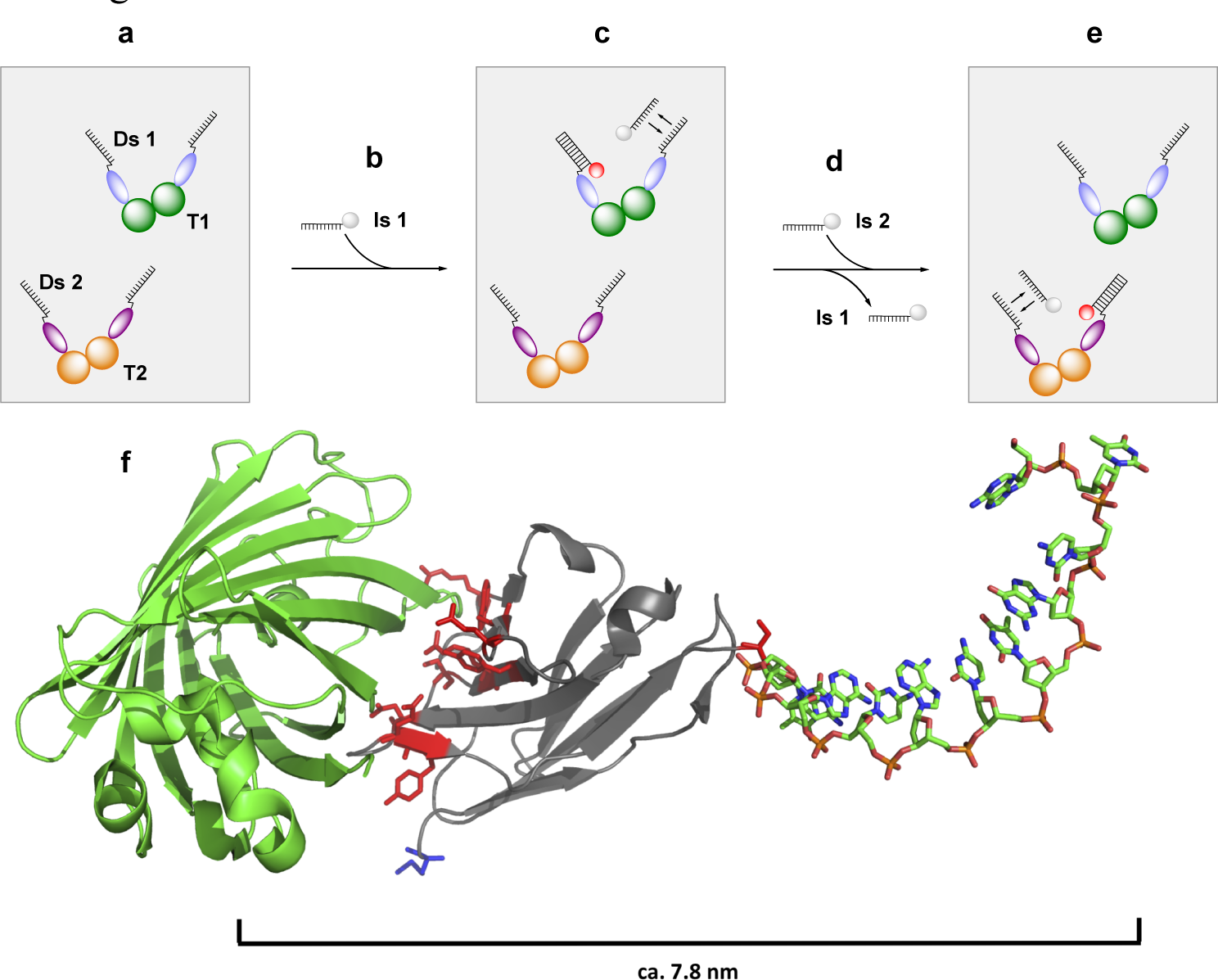
Schematic representation of multicolor DNA-PAINT with two docking-imager strand pairs (modified from [19]). (a) fixed cell stained with two different docking strand (Ds) carrying nanobodies bound to their respective targets. (b) Introduction of fluorophore carrying imager strand 1 (Is1). (c) Image acquisition of target 1. Docking and imager strand transiently interact, bringing the fluorophore into focus and allowing target localization (red). Unbound imager strands freely diffuse and are not localizable (grey). (d) Imager strand 1 is removed by washing and imager strand 2 (Is2) is introduced. (e) Image acquisition and localization of target 2. (f) Model of the docking strand labeled anti-GFP nanobody bound to its GFP target (modified from PDB accession code 3OGO); GFP is shown in green, the nanobody in grey, the nanobody GFP binding residues in red and the nanobody N-terminus in blue. An oligonucleotide is attached C-terminally to illustrate its relative size.

## Methods

### Expression and purification of nanobodies

Expression constructs coding for the anti-tubulin nanobody [22] and anti-GFP nanobody [3,23] were previously described. The anti-tubulin nanobody construct was synthesized by GeneArt Gene Synthesis (Thermo Fisher Scientific) and the anti-GFP nanobody construct was obtained by PCR of pGEX-6P1-containing anti-GFP-nanobody-4K [23]. PCR-amplified constructs were digested with NcoI and Bste l restriction enzymes and ligated in frame in similarly cut pHEN29 plasmid [15]. The pHEN29 plasmid comprises an N-terminal pelB signal sequence, directing the nanobody to the periplasm, followed by a C-terminal Sortase A recognition sequence (LPETGG) upstream of a His_6_ tag and an EPEA tag. Resulting constructs pHEN29-anti-GFP-LPETGG-His_6_-EPEA and pHEN29-anti-tubulin-LPETGG-His_6_-EPEA were confirmed by sequencing and transformed into *E. coli* WK6 for expression. Bacteria were grown in 1 l of Terrific Broth medium (Carl Roth) supplemented with 1 mM MgCl_2_, 0.1% Glucose, 0.4% glycerol and 100 µg/ml ampicillin at 37°C, to an OD_600_ of 0.6 – 0.9 in baffled shaking flasks. Expression was induced with 1 mM Isopropyl-*β*-D-thiogalactopyranoside (IPTG), followed by incubation for 16-18 h at 28°C. Cells were harvested by centrifugation (10 min, 11,800g, 4°C) and the resulting pellets were resuspended in 12 ml ice-cold TES buffer (0.2 M Tris-HCl (pH 8.0), 0.5 mM EDTA, 0.5 M Sucrose) containing one protease inhibitor cocktail tablet (Roche) per 10 ml of TES. After shaking for 1 hour at 4°C, and 200 rpm, 18 ml ice-cold TES/4 (0.05 M Tris (pH 8.0), 0.125 mM EDTA, 0.125 M Sucrose) were added and the cells were further incubated for 1 hour at 4°C. After centrifugation (11,800g, 30 min), the supernatant was collected, and the TES extraction repeated. The resulting periplasmic extracts were pooled and diluted 1:1 in wash buffer (20 mM HEPES (pH 7.5), 300 mM NaCl, 5 mM imidazole 10% glycerol) and applied to a 5 ml HisPur cobalt column, pre-equilibrated in wash buffer at a flowrate of 3 ml per minute using a peristaltic pump. The resin was washed with 10 bed volumes of wash buffer and His-tagged nanobodies were eluted with 20 mM HEPES (pH 7.5), 300 mM NaCl, 500 mM imidazole and 10% glycerol. Protein concentrations in eluted fractions were determined by absorbance at 280 nm using a NanoDrop1000 spectrophotometer (Thermo Fisher Scientific) using the calculated extinction coefficients. The nanobody was further purified and buffer exchanged by size-exclusion chromatography over a Superdex 75 10/300 GL-column (GE Healthcare) in 20 mM HEPES (pH 7.5), 300 mM NaCl and 10% glycerol on an ÄKTA Pure FPLC system. Nanobody-containing fractions were pooled, analyzed by SDS PAGE and concentrated using Amicon concentrators (3000 MWCO).

### Expression and purification of Sortase A

Expression of the SrtA pentamutant (eSrtA, hereafter SrtA) in pET29 (kind gift from David Liu (Addgene plasmid #75144)) was performed essentially as described with minor changes [24]. Briefly, *E. coli* BL21 DE3 were transformed with pET29-Sortase A pentamutant-His_6_ and grown in 1 l of LB medium, supplemented with 50 µg/mL kanamycin at 37°C to an OD_600_ of ~0.5. Expression was induced with 0.5 mM IPTG and the culture incubated at 30°C for 16 hours. Cells were harvested by centrifugation (6000 x g, 4°C, 15 min) and resuspended in 50 ml cold binding buffer (20 mM HEPES pH 7.5, 150 mM NaCl) and centrifuged again. Bacteria were lysed by resuspension in 25 ml ice cold lysis buffer (50 mM HEPES pH 7.5, 300mM NaCl, 5 mM MgCl_2_, 5 mM Imidazole, 10% Glycerol, 1mg/ml DNase I, 1 mg/ml Lysozyme) and sonication on ice. The lysate was cleared by centrifugation (30 minutes, 16000 x g, 4°C) and purified via HisPur cobalt and size-exclusion chromatography as for nanobody purification. SrtA containing fractions were pooled and concentrated with Amicon concentrators (3000 MWCO) to 25 mg/ml in 20 mM HEPES pH 7.5, 300 mM NaCl, 10% Glycerol. The protein was analyzed by reducing SDS PAGE, snap-frozen in 100 µl aliquots in liquid nitrogen and stored at −80°C until further use.

### Sortase A-mediated coupling of clickable moieties (DBCO-amine)

SrtA-mediated functionalization of sortagged nanobodies with Dibenzocyclooctyne-NH_2_ (DBCO-amine) (Sigma-Aldrich # 761540) was carried out by reacting 50 µM nanobody, 150 µM SrtA and 10 mM DBCO-amine (from a 50 mM DMSO stock) in sortase buffer (20 mM HEPES (pH 7.5), 150 mM NaCl, 10 mM CaCl_2_) for 1 hour at 24°C. During the reaction, the GG-His_6_-EPEA peptide was cleaved from the nanobody to form a nanobody:SrtA covalent intermediate linked at the threonine in the LPET-sequence. Subsequent addition of DBCO-amine resolved the intermediate, substituting DBCO for SrtA at the LPET threonine residue. SrtA-His_6_-GG-His_6_-EPEA and unreacted His-tagged nanobody were separated from DBCO-nanobody conjugate using HisPur cobalt resin. For this, the reaction mixture was diluted 1:1 in 20 mM HEPES (pH 7.5), 300 mM NaCl, 10% Glycerol, 5 mM Imidazole, supplemented with 10 mM EDTA to stop the sortase reaction. Resin was washed with wash buffer until UV absorption at 280 nm reached baseline. Nanobody-DBCO containing flowthrough and wash fractions were pooled and remaining DBCO-amine was removed using an Amicon concentrator (3000 MWCO). Nanobody-DBCO-conjugate concentrations were determined by Bradford-assay and successful conjugation and purity were assessed by SDS-PAGE. The degree of labeling (DOL) was determined by absorbance at 280 nm (nanobody) and 309 nm (DBCO) using a Nanodrop1000 spectrophotometer (Thermo Fisher Scientific) (Supplementary Table 2). The conjugate was stored at 4°C or immediately used for click-reactions. Optimal molar ratios of DBCO-amine:nanobody and reaction duration were determined in test reactions with 50 M nanobody with 150 M SrtA pentamutant in sortase buffer at 24°C, with a DBCO-amine concentration between 2.5 and 12.5 mM for 0.5-5 hours.

### Copper-free SPAAC with azide-reaction partner

Docking strand-oligonucleotides (Binder 1: TTATACATCTAG and Binder 3: TTTCTTCATTA), modified with a 5’ azide moiety were synthesized by Microsynth (Switzerland) and dissolved in PBS (pH 7.4) (Sigma Aldrich) to a concentration of 10 mM. Alexa Fluor 647 (Invitrogen) containing an azide moiety was dissolved in DMSO to a final concentration of 10 mM. For a standard reaction, 40 µM DBCO-nanobody conjugate was mixed with 40 µM azide-reaction partner in 20 mM HEPES (pH 7.5), 300 mM NaCl and 10% glycerol and incubated for 1 h. Unbound reaction partners were removed in two buffer exchange steps with a Zeba spin desalting column (7000 MWCO). Successful labeling was verified by reducing SDS PAGE and analysis on a FUSION-FX7 Spectra fluorescence imager (Vilber Lourmat). The DOL was determined as for DBCO, using absorbance at 650 nm for the dye (Supplementary Table 2). Optimal molar ratios for the reaction were determined using azide-modified methoxy-polyethyleneglycol (mPEG, M_W_ = 2000) as a click-reaction partner in molar ratios of nanobody to azide-PEG ranging from 1:1 to 1:10 and incubating for 0.5 to 12 hours. Reactions were quenched by adding 1 mM DBCO-amine to the mixtures. Samples were analyzed on reducing SDS PAGE.

### Cell culture and immunostaining

HeLa cells, stably expressing CAV1-GFP (kind gift of the Helenius laboratory [25]) were maintained in DMEM (Life Tech) supplemented with 10% FCS and 1% Glutamax (Life Tech) at 37°C and 5% CO_2_ in a humidified incubator. After 24 hours, cells were washed 3x with PBS and incubated in pre-extraction buffer (80 mM PIPES (pH 6.9), 1 mM MgCl_2_, 10 mM EGTA, 0.5 % Triton X-100) for 60 seconds at room temperature. Pre-extraction buffer was replaced by fixation buffer (80 mM PIPES (pH 6.9), 1 mM MgCl_2_, 1 mM EGTA, 3.2 % paraformaldehyde, 0.1 % glutaraldehyde) and cells were incubated for 10 minutes at 37 °C. Fixation buffer was removed by washing with PBS and 10 mM freshly prepared sodium borohydride in PBS was then added for 7 minutes followed by a ten-minute incubation in 100 mM glycine. Fixed cells were repeatedly washed with PBS and blocked with Image iTFX signal enhancer (Invitrogen) for 1 hour and then with 5 % BSA and 0.1 % Triton X-100 in PBS for 30 min at room temperature. After washing 3x with PBS, cells were stained with a 2.5 µg/ml solution of nanobodies, diluted in 5 % BSA and 0.1% Triton X-100 in PBS. After 1 hour at room temperature, cells were washed 3x with PBS and immediately used for imaging.

### Optical setup

Images were acquired with a Vutara 352 super resolution microscope (Bruker) equipped with a Hamamatsu ORCA Flash4.0 sCMOS for super resolution imaging and a 60x oil immersion TIRF objective with numerical aperture 1.49 (Olympus).

### Super-resolution dual color DNA PAINT imaging

Imager strand oligonucleotides (Imager 1: CTAGATGTAT and Imager 3: GTAATGAAGA), modified with a 3’ Atto 655 dye, were synthesized by Eurofins Genomics (Germany), aliqoted in PBS at a concentration of 100 µM and stored at −20 °C until imaging. For each sample, imager strand concentrations were empirically determined to ensure sufficiently high on-off ratios. Fixed samples were placed in a one-well magnetic chamber (Live Cell Instrument, South Korea) and covered in PBS (pH 7.4) supplemented with 0.5 M NaCl and 500 pM Atto 655-labeled Imager 1. A 250 pM solution of Atto 655 labeled Imager 3 was then added. Ideal exposure times for both binder-imager pairs were calculated with the Picasso Render tool [26]. Data were acquired with TIRF/HILO-illumination at a laser-power density of 2.5 kW/cm^2^ using a 639 nm laser. Images were collected with a 100 ms acquisition time and typically 10000 images were used to reconstruct the super-resolution composites.

### Super-resolution data processing

Raw data files were analyzed using the Picasso software package (https://github.com/jungmannlab/picasso). Individual binding events were identified by Picasso:Localize (Identification) with a box side length of 7 and a minimum net gradient of 1500 and 2000 for the microtubule and caveolae localizations, respectively. Subsequently, the localizations were loaded into Picasso:Render to correct for microscopic drift (Undrift by RCC), align microtubule and caveolin fluorescence and obtain binding kinetics. For our complementary oligos, we determined a t_on_ = 600 ms and t_off_= 118 s for binder-imager pair 1 and t_on_ = 520 ms and t_off_= 153 s for binder-imager pair 3. Microtubule (250 nm straight sections without overlaps, aligned and processed as in [8]) and caveolin cross sections were taken in ImageJ and fitted to a Gaussian function to determine their respective full width at half maximum and interpeak distance.

## Results

### Nanobody production

Recombinant anti-tubulin and anti-GFP-LPETGG-His_6_-EPEA nanobody constructs were produced in *E. coli*, extracted from the bacterial periplasm via osmotic shock and purified via immobilized metal ion and size exclusion chromatography to near homogeneity (>95% by SDS-PAGE). Typical final yields were between 15 and 20 mg/l culture for both constructs (Supplementary Figure 1). SrtA (ca. 40 mg/l culture) was similarly produced and purified.

### SrtA-mediated conjugation and copper-free SPAAC via DBCO-amine

For derivatizing nanobodies with short oligonucleotides to use as labels in DNA-PAINT, we initially attempted a peptide exchange strategy using SrtA, similar to that of Massa *et al.* [15]. We used a SrtA pentamutant [24] to label two different probes: an anti-tubulin [22], and an anti-GFP nanobody [27]. Our substrate peptide carried a C-terminal alkyne moiety providing a specific reaction point for azide labeled ‘docking’ oligonucleotides using click chemistry via Cu(I) catalysed alkyne-azide cycloaddition (CuAAC) [28]. Although the reactions were successful (data not shown), we encountered several difficulties with the procedure (see Discussion), not least of which was the poor yield of final labeled nanobodies due to the need to remove Cu(I) from the reaction mix. In order to simplify the labeling procedure and maximize the yield, we sought to devise an efficient Cu(I) independent reaction scheme. Cu(I) free click reactions with biomolecules based on strain promoted alkyne-azide cycloaddition (SPAAC) using dibenzocyclooctyne (DBCO) have been reported and can be performed under physiological conditions [28–30]. Building on this idea, we tested whether an amine modified DBCO could directly function as the SrtA releasing nucleophile (Figure 2a). We again prepared SrtA-nanobody intermediates and subsequently added DBCO-amine in excess. We observed nearly quantitative conversion of the SrtA-nanobody intermediate and confirmed the association of one DBCO molecule per nanobody by absorption spectroscopy (Figure 2c, f; Supplementary Table 2) and Bradford assay. To confirm that the resulting constructs were competent for label addition, we optimized the labeling reaction using azide-modified mPEG (M_W_ = 2000 Da) at different concentrations and varying incubation times and monitored the results by electrophoretic mobility shift on SDS polyacrylamide gels. Nearly quantitative labeling was observed after 12 hours incubation with a 1:1 molar ratio of azide-modified reaction partner (Supplementary Figure 2). To assess whether the reaction yields conjugates with only one label per protein, Alexa-647-azide was reacted with nanobody:DBCO. Addition of Alexa-647-azide to the DBCO-nanobodies in a 1:1 ratio resulted in association of 1 fluorophore per nanobody, again confirmed by absorption spectroscopy (Figure 2d, e; Supplementary Table 2). The nanobodies performed well in conventional (d)STORM SMLM (not shown).

**Figure 2.**
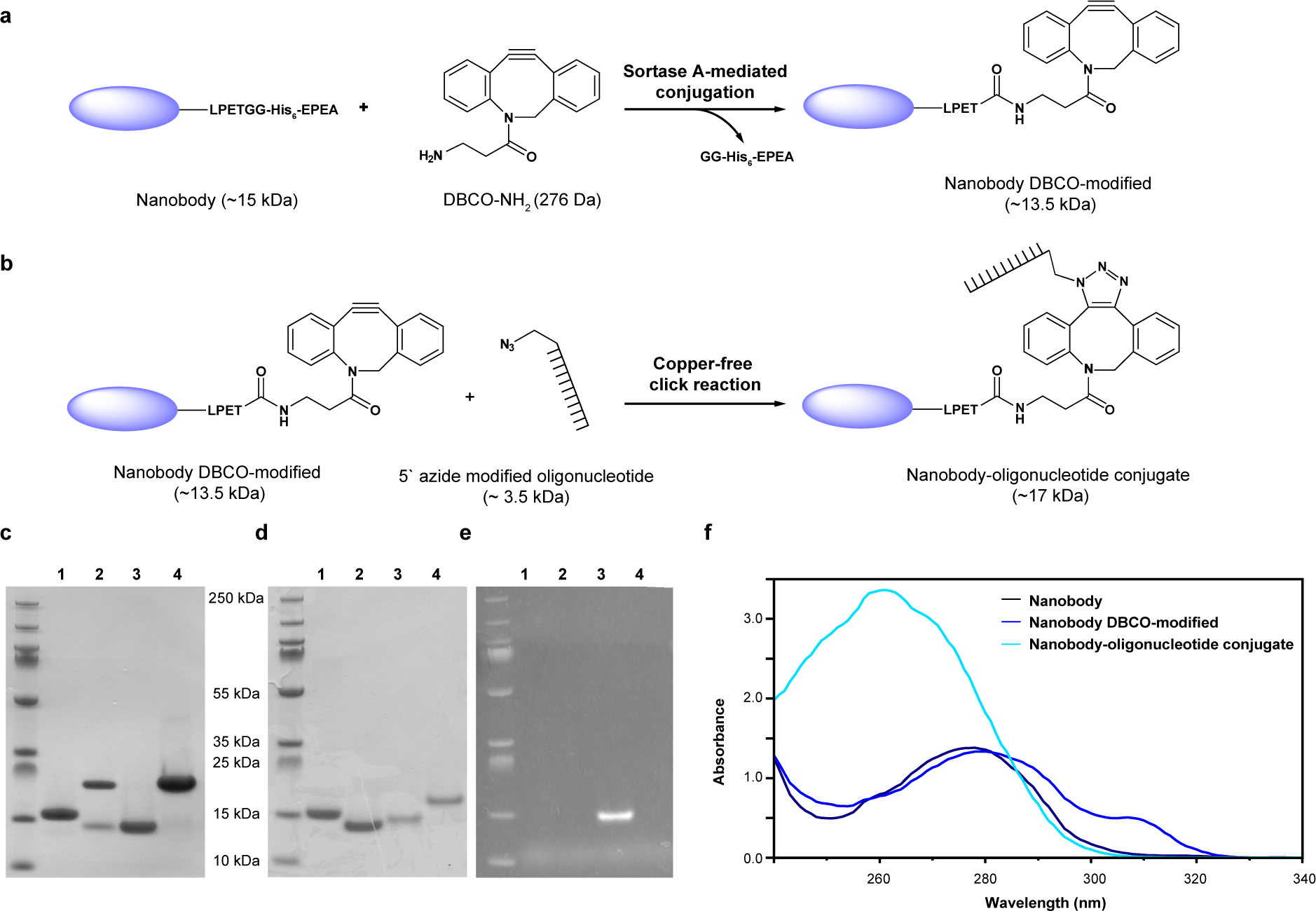
Schematic representation of the site-specific labeling of nanobodies via SrtA. (a) Srt A-mediated conjugation of DBCO amine to sortagged nanobody. (b) Copper-free click reaction of the nanobody-DBCO conjugate with azide-modified oligonucleotide. (c) Coomassie-stained SDS PAGE gel of samples taken during the SrtA-mediated conjugation of DBCO amine to anti-GFP nanobody. 1 = unmodified anti-GFP nanobody; 2 = reaction mixture after 1 h incubation; 3 = pooled and concentrated DBCO-modified anti-GFP nanobody; 4 = elution of remaining His-tagged anti-GFP nanobody and SrtA. (d) Coomassie-stained SDS PAGE gel of samples taken before and after the click reaction. 1 = unmodified anti-GFP nanobody; 2 = DBCO modified anti-GFP nanobody; 3 = Alexa Fluor 647 conjugated anti-GFP-nanobody; 4 = docking strand 3 conjugated anti-GFP nanobody. (e) Fluorescence image of the SDS PAGE gel shown in (e), confirming fluorophore association. (f) UV absorbance spectra of the different conjugates.

We then repeated our labeling reaction using azide-oligo docking strands. Again, we observed high efficiency addition of the oligos to the nanobodies based on electrophoretic mobility shift on SDS gels (Figure 2b, c).

### Dual color PAINT labeling

The anti-tubulin- and anti-GFP-LPETGG-His_6_-EPEA nanobody constructs were coupled to previously reported oligo sequences binder 3 and binder 1, respectively [16]. The high efficiency of the SPAAC reaction of DBCO-functionalized nanobodies with the azide-modified docking strands allowed easy removal of unreacted oligonucleotides by simple buffer exchange.

We tested the viability of our nanobodies as super-resolution labeling probes by staining fixed Hela cells stably expressing CAV1-GFP with anti-tubulin nanobody coupled to binder 3 and anti-GFP nanobody coupled to binder 1. Sequential addition of dye-labeled imager 3 and imager 1 strands allowed us to acquire dual color DNA-PAINT images of cells (Figure 3). The microtubules were resolved with a full width half maximum of 31 ± 4 nm and the plasma membrane caveolae with a diameter of 61± 17 nm (Supplementary Figures 4, 5). These distances are far lower than the diffraction limit of light and agree well with previously reported microtubule [22] and caveolae diameters *[31]*.

**Figure 3.**
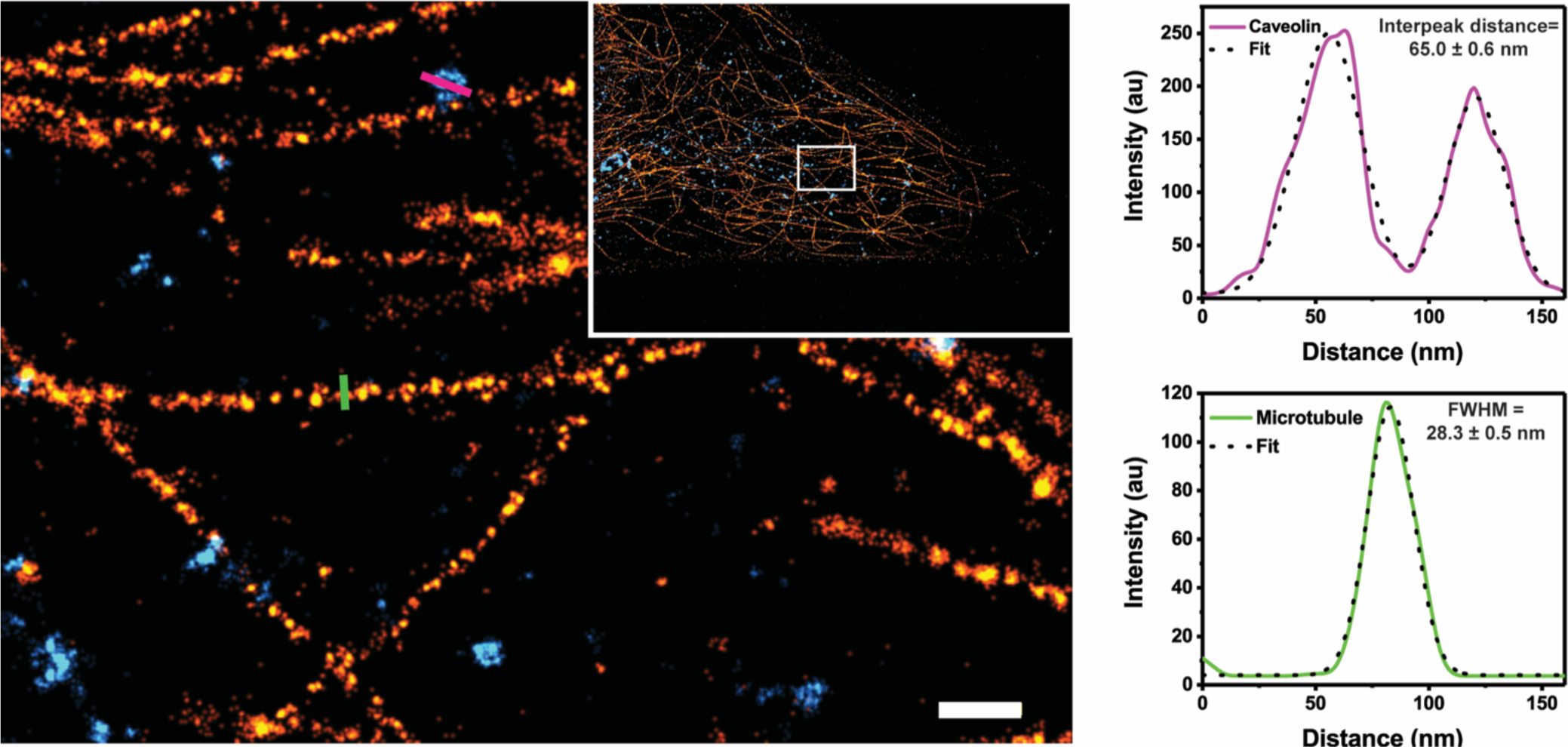
Dual color DNA-PAINT image of tubulin (orange) and caveolin (blue). On the left, resolved microtubules labeled with anti-tubulin-nanobody and caveolin-GFP labeled with anti-GFP-nanobody. Scale bar is 100 nm. Inset, overview of the whole cell. The enlarged area is indicated with a white box. On the right, cross sections of a typical caveola (magenta) and microtuble (green) with full width at half maximum (FWHM) given for each, along with the fitting error. The NeNa localization precision [32] was 9 nm for both the microtubule and caveolin localizations.

## Discussion

The efficient conjugation of labels to an antigen-specific antibody or nanobody is a crucial first step for super resolution microscopy studies. Conventional protocols rely primarily on the use of N-hydroxysuccinmide (NHS)-mediated conjugation of dyes to the ε-amine groups of lysine side chains. Since most proteins harbor several solvent-exposed lysines, this precludes site-specific label introduction and leads to non-uniform labeling of the probe, although specific labeling of the N-terminus has been reported [19]. Lysine residues can also occur near the antigen-binding site in the labeled protein, which can result in loss of affinity or complete inhibition of binding the target after dye conjugation.

Introducing an unpaired carboxy-terminal cysteine for conjugation via maleimide chemistry allows the site-specific introduction of a label without disturbing binding performance [33], but since IgGs and nanobodies require disulfide bonds for stability, the introduced cysteine can interfere with their proper folding. Even if this is not the case, the necessary reduction step prior to the labeling reaction can cause a substantial loss of material due to inadvertent reduction of the obligate disulfides [15]. Other site-specific methods, including SrtA-mediated labelling, make use of chemoenzymatic reactions whose exquisite specificity guarantees the stability and homogeneity of labeling. This is especially desirable for DNA-PAINT labeling, as only stoichiometric labeling will make the probe compatible with quantitative approaches like qPAINT [34]. Since the oligonucleotide is quite large in comparison with the nanobody (see Figure 1f), its coupling close to the epitope-binding site in combination with its high charge may significantly perturb target recognition.

The SrtA variant from *Staphyloccocus aureus* recognizes an LPXTG-sequence (where X = any amino acid) on the target protein which it cleaves at the terminal glycine to form a target-SrtA acyl-intermediate at the preceding threonine, simultaneously releasing the C-terminal fragment. Subsequent nucleophilic attack by an N-terminal amine-containing species then releases SrtA, generating a hybrid target. The reaction is exploited for labeling by using probes containing a primary amine that can resolve the target-SrtA intermediate [35–37]. This reaction scheme allows specific coupling of a wide range of functional molecules to the C-terminus of any LPXTG-containing protein, including ‘clickable’ alkyne residues.

While CuAAC has been successfully applied for a variety of biomolecules, it requires a multi-step protocol and has the disadvantage of Cu(I) toxicity, thereby requiring removal of the metal for live cell experiments; additionally, Cu(I) can have a denaturing effect on proteins. Due to the need to remove Cu(I) from our reaction mixtures, we found it difficult to obtain satisfying yields of labeled nanobody. We therefore sought to eliminate Cu(I) altogether and reduce the number of steps in the labeling procedure by using SPAAC.

Van Lith *et al.* [38] used dibenzocyclooctyne (DBCO) in SPAAC as a general reactive moiety attached to a variable length PEG amine to provide a functionalizable linker for derivatization using various reaction chemistries. While this flexibility in linker length is intended to be beneficial for preparing antibody-drug conjugates (ADCs) for *in vivo* applications, it confers no advantage for super resolution fluorescence studies, where increasing linker length complicates the precise localization of the detected species. We therefore used DBCO-amine directly.

Baer *et al*., [39] in testing different amine nucleophiles for resolving the Srt A intermediate, observed a clear preference for small amine groups for the most efficient reactions. In our hands, resolution of the SrtA reaction using DBCO-amine was nearly quantitative. The specificity and low cost of the reagent provides the additional benefit that it can be used in large excess in the reaction to out-compete other potential nucleophiles (including the initially cleaved NH2-GGG-His6 peptide), thereby driving the desired reaction to completion.

By using DBCO-amine instead of Cu(I) in the reaction, we substantially reduced the preparation time of our samples while simultaneously increasing the efficiency of the labeling reaction and the overall yield of labeled probe. We suggest that the results here can be extended to accommodate the most diverse labeling strategies by utilizing small, bifunctional molecules having any compatible reactive group of interest coupled to an amine moiety that can complete the Sortase reaction. This approach facilitates the site-specific introduction of uniquely reactive tags into any peptide of interest, the hydrodynamic radius of whose subsequent reaction products need not exceed what the label itself adds. As the native Sortase reaction requires a small amine nucleophile (typically an N-terminal glycine for peptides), the high yields and simplified preparation of our nanobody conjugates achieved using DBCO-amine are likely to be mimicked using other small amine nucleophiles, greatly improving the ease and flexibility of preparing probes for super resolution studies. In our case, addition of the detection moiety to the C-terminus of the nanobodies ensures an accessible recognition sequence and that the modification has no detrimental effect on the binding interface, making the final constructs – always with a single label per molecule - ideal for a variety of quantitative studies (for example, single molecule brightness analyses, stepwise photobleaching, etc.). Taken together we here present a simple and efficient method for quantitative labeling of nanobodies for DNA-PAINT that due to the versatility of copper-free click chemistry provides an available labeling site at the very C-terminus of nanobodies for a variety of labeling schemes.

One of the great advantages of DNA-PAINT is the possibility of multiplexed labeling with the same dye on different epitopes where the specificity is encoded in the DNA-oligomers [16]. The combination with nanobodies holds great promise as more and more such binders are developed. In the future, highly multiplexed labeling will be possible using secondary nanobodies [40], nanobodies against peptide tags [41] and RFPs [14]. Further optimization of the labeling procedure will likely allow for unprecedented resolution using DNA-PAINT in cells.

## Acknowledgements

We thank Jan Gettemans and Ralf Jungmann for advice and all members of the Ewers laboratory for helpful discussions. This work was supported by DFG through SFB958 INST 130/827-2 to H.E.

---

**Supplementary Table 1:** Calculated molecular weight (MW) of nanobodies throughout the conjugation process.

|  | anti-tubulin nanobody | anti-GFP nanobody |
| --- | --- | --- |
| MW of nanobody-LPETGG-His<sub>6</sub>-EPEA<sup>1</sup> | 15060 Da | 14591 Da |
| MW of nanobody-LPET<sup>1</sup> | 13697 Da | 13227 Da |
| MW of nanobody-LPET-DBCO<br>(MW<sub>DBCO amine</sub> ≈ 276 Da) | 13973 Da | 13503 Da |
| MW of nanobody-DBCO-mPEG<br>(MW<sub>mPEG</sub> ≈ 2000 Da) | 15973 Da | 15503 Da |
| MW of nanobody-DBCO-AlexaFluor 647<br>(MW<sub>AF647</sub> ≈ 850 Da) | 14823 Da | 14353 Da |
| MW of nanobody-DBCO-docking strand oligo<br>(MW<sub>oligo</sub> ≈ 3700 Da) | 17673 Da | 17203 Da |

<sup>1</sup> Calculated with ExPASy ProtParam tool (expasy.org/protparam/)

**Supplementary Table 2:** Yield and degree of labelling (DOL) of SrtA-mediated labelling procedure and SPAAC reaction. Values were calculated from measurements of three independent labeling reactions. Errors indicate the standard deviation of the averaged values.

|  | anti-tubulin nanobody | anti-GFP nanobody |
| --- | --- | --- |
| Total protein output/ input of the SrtA reaction x 100%<sup>2</sup> | 90 % | 93 % |
| Average number of DBCO functionalities incorporated per nanobody (DOL<sub>DBCO</sub>)<sup>3</sup> | 0.91 ± 0.06 | 0.90 ± 0.04 |
| Average number of AlexaFluor 647 azide labels per nanobody (DOL<sub>AF647</sub>)<sup>3</sup> | 0.94 ± 0.03 | 0.93 ± 0.05 |

<sup>2</sup> Total protein concentration of nanobody used for the reaction (input) and nanobody after the SrtA-mediated conjugation and purification (output) were assessed with Bradford protein assay.
<sup>3</sup> Absorbance of the nanobodies ( $A_{280}$ ) and the labelling probe ( $A_{max}$ ) were measured spectrophotometrically. DOL was calculated using Formula (1) with the calculated extinction coefficients of the respective nanobody ( $\epsilon_{nanobody}$ ), of the labelling probe ( $\epsilon_{max}$ ) and the correction factor for the absorbance of the labelling probe at 280 nm (CF).
With $\epsilon_{anti-GFP-LPET} = 27,055 \text{ M}^{-1} \text{ cm}^{-1}$ ; $\epsilon_{anti-tubulin-LPET} = 35,535 \text{ M}^{-1} \text{ cm}^{-1}$ ; $\epsilon_{max}(\text{DBCO}) = 12,000 \text{ M}^{-1} \text{ cm}^{-1}$ ; $\epsilon_{max}(\text{AF647}) = 270,000 \text{ M}^{-1} \text{ cm}^{-1}$ ; $CF_{\text{DBCO}} = 1.089$ ; $CF_{\text{AF647}} = 0.03$ ; $\lambda_{max}(\text{DBCO}) = 309 \text{ nm}$ ; $\lambda_{max}(\text{AF647}) = 650 \text{ nm}$

**Supplementary Figure 1.**
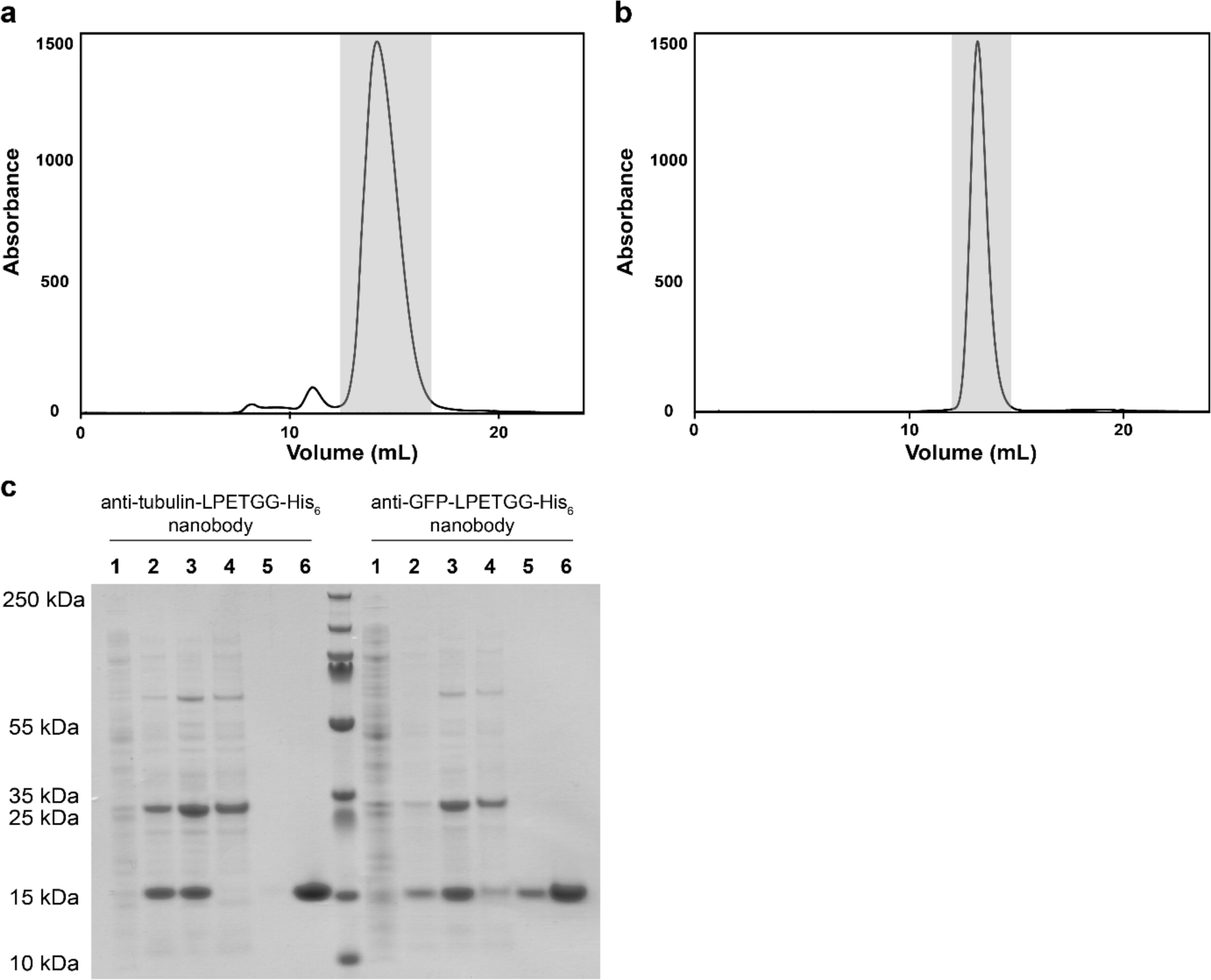
Expression of anti-tubulin and anti-GFP nanobody-LPETGG-His_6_-EPEA constructs. (a) FPLC size-exclusion chromatogram of anti-tubulin nanobody-LPETGG-His_6_-EPEA elution from HisPur resin. Nanobody-containing fractions are highlighted in grey. (b) FPLC chromatogram of anti-GFP nanobody-LPETGG-His_6_-EPEA elution from HisPur resin. Nanobody-containing fractions are highlighted in grey. (c) Coomassie-stained SDS PAGE gel of samples taken during the expression and purification of nanobodies. Lane 1 = induced expression culture; lane 2 = TES extract I; lane 3 = TES extract II; lane 4 = HisPur cobalt purification flow-through; lane 5 = HisPur cobalt purification pre-elution; line 6 = pooled nanobody-containing fractions from the peaks highlighted in grey in (a) and (b).

**Supplementary Figure 2.**
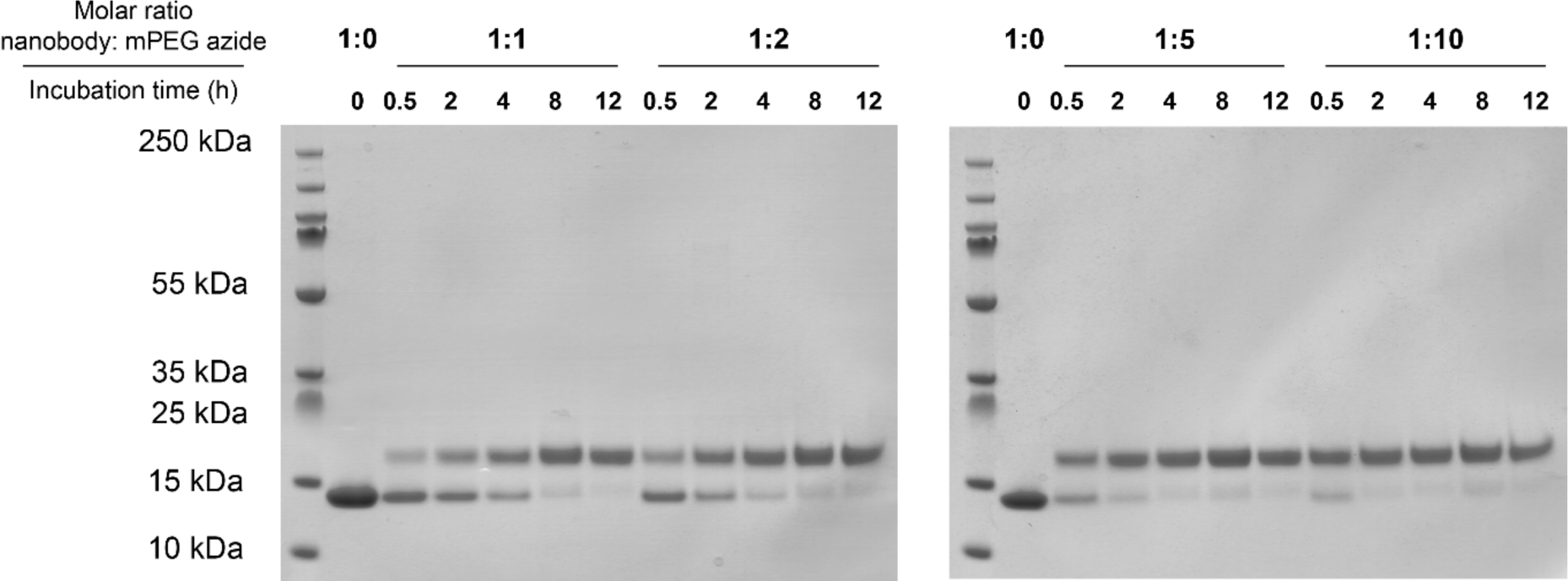
Reducing SDS PAGE gels of the optimization of molar ratio and incubation time of the click reaction between nanobody-DBCO-conjugate and azide reaction partner, using mPEG-azide (MW = 2000 Da) and anti-GFP-DBCO nanobody conjugate.

**Supplementary Figure 3.**
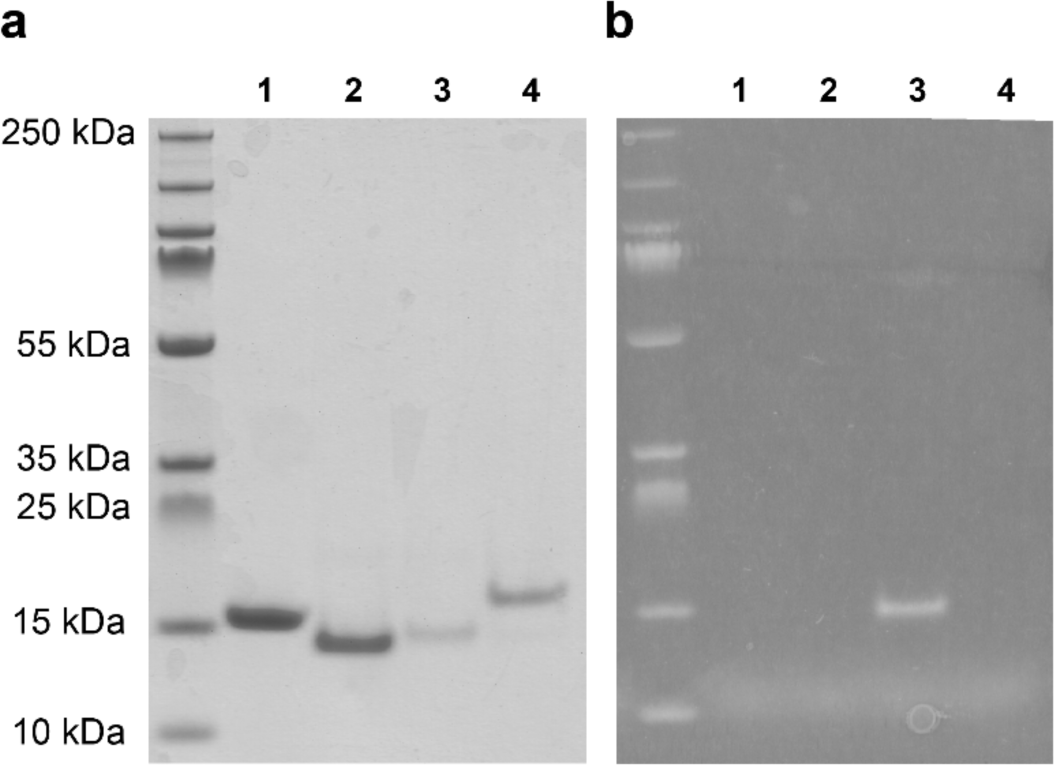
Analytical SDS-PAGE of click reaction. (a) Coomassie-stained reducing SDS gel of samples taken before and after the click reaction. Lane 1 = unmodified anti-tubulin nanobody; lane 2 = DBCO modified anti-tubulin nanobody; lane 3 = Alexa Fluor 647 conjugated anti-tubulin-nanobody; lane 4 = docking strand 3 conjugated anti-tubulin nanobody. (b) Fluorescence image of the SDS PAGE gel shown in (e), confirming fluorophore association.

**Supplementary Figure 4.**
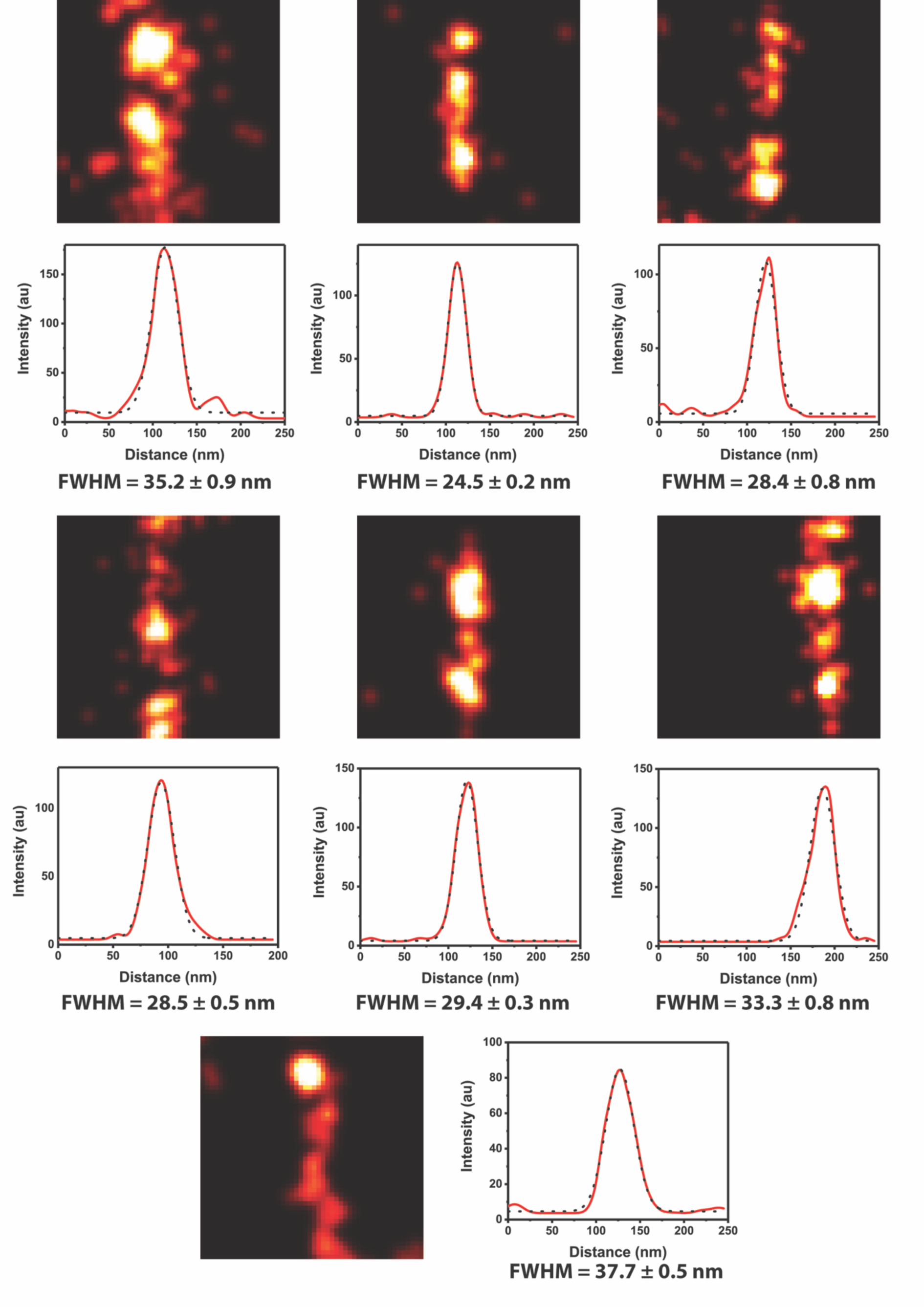
Cross-sections of 7 different microtubule segments. Straight microtubule sections from 7 different areas (250 × 250 nm) of the super-resolved images have been taken. Their cross-section is plotted and fitted with a Gaussian, from which the full-width half maximum (FWHM) was calculated. The error represents the fitting error. Taken together, the microtubules had a mean FWHM of 31 ± 4 (standard deviation) nm.

**Supplementary Figure 5.**
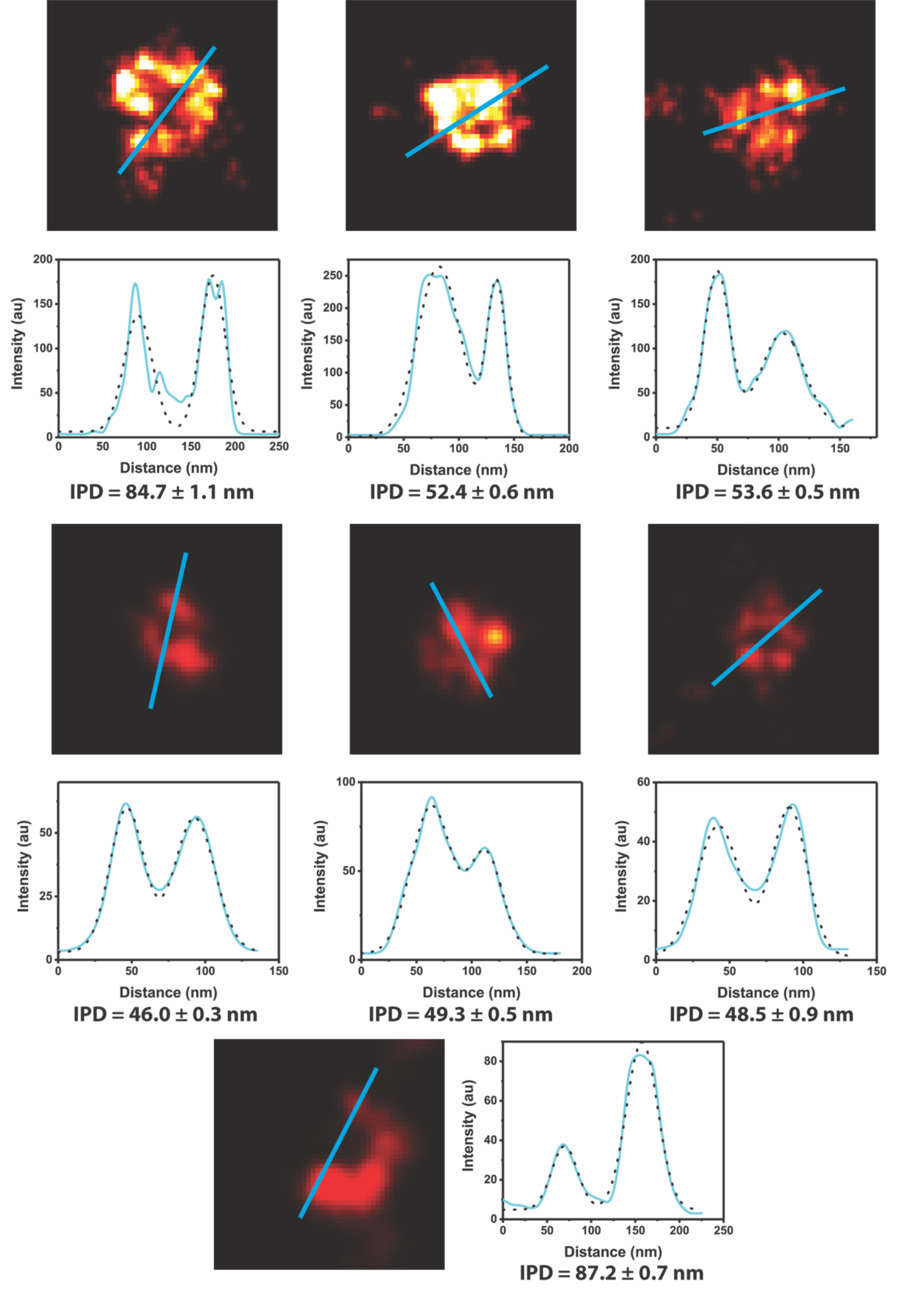
Cross-sections of 7 different caveolae. From the super-resolved images, 7 different caveolae have been picked (250 × 250 nm areas), their cross-section was plotted and fitted with a double Gaussian. The interpeak-distance (IPD) was calculated with the error representing the fitting error. Taken together, the caveolae had a mean diameter of 61 ± 17 (standard deviation) nm.

